# Identification and study of Prolyl Oligopeptidases and related sequences in bacterial lineages

**DOI:** 10.1101/2024.09.22.614393

**Authors:** Soumya Nayak, Ramanathan Sowdhamini

## Abstract

Proteases are enzymes that break down proteins, and serine proteases are an important subset of these enzymes. Prolyl oligopeptidase (POP) is a family of serine proteases (S9 family) that has the ability to cleave peptide bonds involving proline residues and it is unique for its ability to cleave various small oligopeptides shorter than 30 amino acids. The S9 family from the MEROPS database, is classified into four subfamilies based on active site motifs. These S9 subfamilies assume a crucial position owing to their diverse biological roles and potential therapeutic applications in various diseases. In this study, we have examined ∼32000 completely annotated bacterial genomes from the NCBI RefSeq Assembly database to identify annotated S9 family proteins. This results in the discovery of ∼53,000 bacterial S9 family proteins (referred to as POP homologues). These sequences are classified into distinct subfamilies through various machine-learning approaches and comprehensive analysis of their distribution across various phyla and species and domain architecture analysis are also conducted. Distinct subclusters and class-specific motifs of POPs were identified, suggesting differences in substrate specificity in POP homologues. This study can enable future research of these gene families that are involved in many important biological processes.

## Introduction

A protease is an enzyme that facilitates the breakdown of proteins into smaller peptides or amino acids by catalysing the hydrolysis of peptide bonds. Proteases are involved in a wide range of biological processes including digestion, cellular signalling, and immune response. There are several classes of proteases including serine proteases, cysteine proteases, aspartate proteases, and threonine proteases. Each class of protease has its own characteristic mechanism of action and substrate specificity.

Serine proteases catalyse the hydrolysis of peptide bonds in proteins. Their active site is characterized by the presence of a conserved active site Serine, which acts as a nucleophile during the catalytic reaction. In fact, about one-third of the proteases can be classified as serine proteases (1). More than 50 families of Serine Protease have been classified by the MEROPS (2) database.

These include subtilisin-like, chymotrypsin-like, and trypsin-like proteases. They function in diverse biological processes such as digestion, blood clotting, fertilisation, development, immune response, and secondary metabolism (3)(4). Serine proteases in prokaryotic lineages are diverse enzymes with critical functions and are involved in several functions associated with cell signalling, defence response and development (5)(6). Their activities are tightly regulated to maintain cellular homeostasis and contribute to the survival and adaptability of organisms in various environments. Prolyl oligopeptidase (POP) is a serine protease enzyme that specifically cleaves peptide bonds involving proline residues. Prolyl oligopeptidase is a member of the S9A family according to the MEROPS database. It is a distinct serine protease that exhibits the ability to hydrolyse internal proline residues. Unlike other proteases, POP can cleave peptide bonds involving proline due to its ability to accommodate the imino ring structure of proline residues. This unique feature allows POP to selectively target oligopeptides of up to 30 amino acids in length (7). This is different from other proteases which primarily act on bigger proteins. POPs are found across various domains of life, including bacteria, archaea, and humans, highlighting their widespread distribution and evolutionary significance (8).

The hallmark of POPs is the presence of a catalytic α/β hydrolase domain and a β-propeller domain. POPs are typically around 700 residues in length in most species (7). Studies on the crystal structure of bacterial POPs(S9A) from *Myxococcus xanthus, Sphingomonas capsulata*, and *Aeromonas punctata* have revealed a two-domain architecture (9)(10), similar to mammalian POPs. The α/β hydrolase domain in POPs is characterized by a short helical N-terminal region (∼70 residues) and a larger C-terminal region that encompasses the catalytic triad. This catalytic region is sequentially very less conserved to other α/β hydrolases, but structurally highly superimposable. Notably, the catalytic triad composed of Ser, Asp, and His residues is concealed at the interface between the two structural domains (Fig 1A).

**Fig 1.**
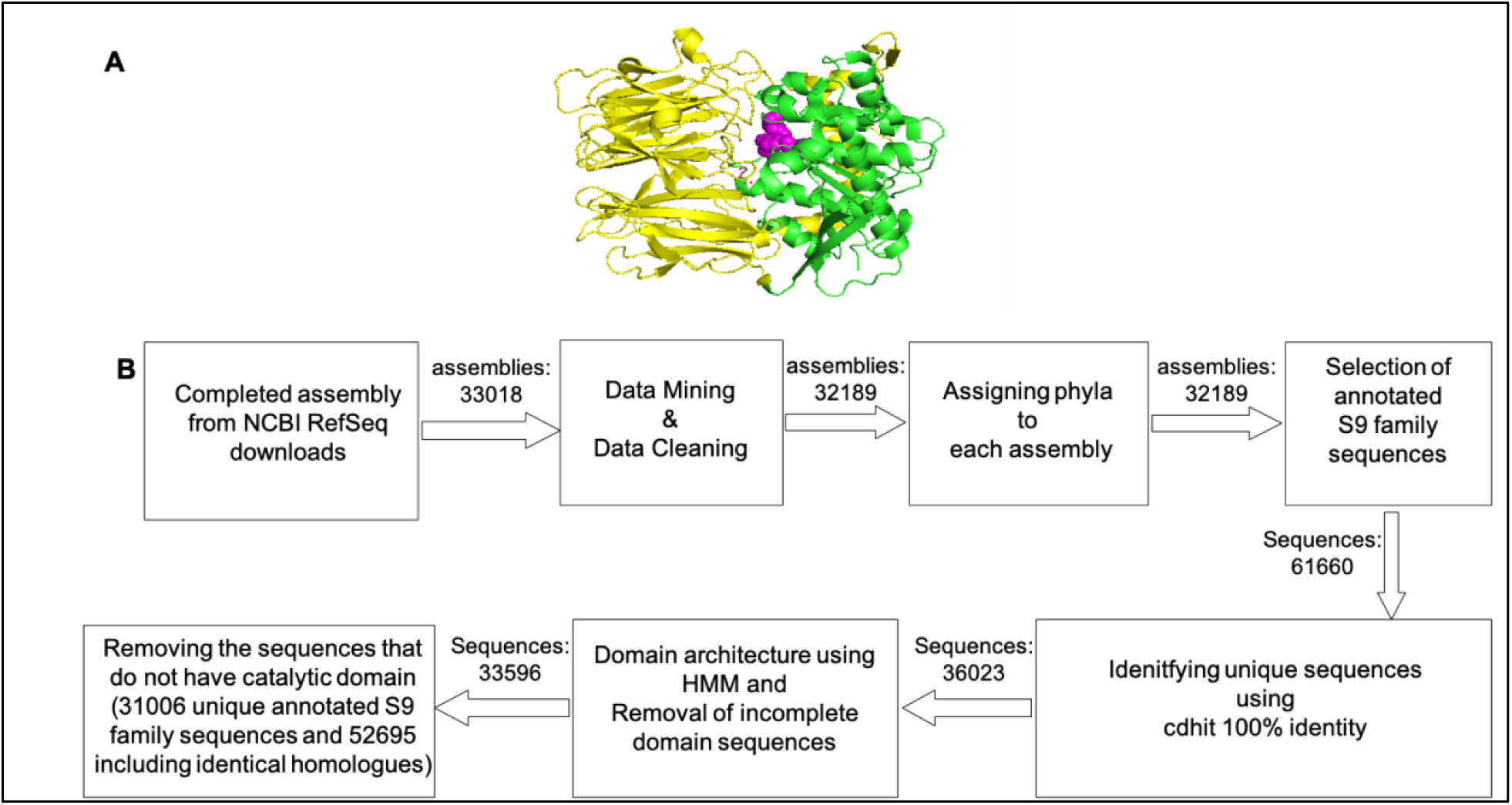
Pipeline: Fig 1A represents a representative POP structure (pdb structure **3ivm**). Catalytic domain(green) and propeller domain(yellow) are shown. The active site residues are marked in magenta. The pipeline followed in the study is highlighted in Fig 1B.

According to the MEROPS database, the S9 family is divided into four subcategories (11) S9A, S9B, S9C and S9D. Different members have different substrate specificity. The active site residues are in the order Ser, Asp, and His in the primary sequence. The motifs around the active site, Serine, are characteristic for each subfamily: GGSXGGLL, where X is typically Asn or Ala (for S9A), GWSYGGY (for S9B), GGSYGG (for S9C) and GGHSYGAFMT (for S9D) (11). POPs(S9A) cleave peptide bonds at the C-terminal side of proline residues. OPBs (S9A) cleaves Arg+ and Lys+ bonds. DPP IV(S9B) cleaves dipeptides where the penultimate amino acid is proline (12). DPPs are homodimers and exist in both soluble and membrane-bound forms whereas POPs and OPBs are monomers. Aminoacyl peptidase(S9C) cleaves enzymes from the blocked amino terminal residues of peptide (13). Glutamyl endopeptidase cleaves the synthetic substrate Z-Leu-Leu-Glu-naphthylamide.

Classification of the sequences is important as the different members of the superfamily have different biological significance. Yet sequences mostly are annotated as S9 family proteases. POPs are targeted as therapeutic for celiac disease as well as implicated in neurological disorders (14)(15). OPBs are considered as a potential drug target for Trypanosoma disease (16). DPPs of clinical isolates of *P. Gingivalis* strains with high dipeptidyl peptidase 4 (DPP4) expression also have a high capacity for biofilm formation and were more infective (17). The role of bacterial DPP-IV in human diseases is still emerging, and further research is needed to explore its implications. In addition, DPP IV is responsible for the degradation of incretins, such as GLP-1 (18) and plays a major role in glucose metabolism. Thus, DPP IV inhibitors are a well-known target of a new class of oral hypoglycaemics (19). Some research implies that DPP4 from gut bacteria can also decrease the active glucagon-like peptide-1 (GLP-1) and disrupt glucose metabolism (20). There is also evidence that bacterial acyl aminoacyl peptidase is used by certain bacteria for cold adaptation (21).

Previous analysis from our lab had considered 1,202 bacterial and 91 archaeal genomes and revealed ∼3,000 POP homologues. Out of 3000 homologues, only 638 were annotated as POPs. A significant extension of this gene family was also proposed by characterizing 39 new POPs and 158 new α/β hydrolase members (22). Since this analysis, the NCBI assembly repository has increased at least 30-fold. We have now considered ∼32000 bacterial genomes, which have complete annotations in the NCBI assembly database, for the current study. The availability of such a large genomic/proteomic information of many bacterial species offers a great opportunity to understand the detailed distribution, classification domain architecture and other chemical/biological properties of Bacterial S9 family peptidases. In this study, we presented the underlying preliminary nature of the annotation of the S9 family members in the bacterial genomes. We also explained the need for a machine-learning tool for enhanced functional annotation for more specific sub-category annotation and also provided an ML solution based on Protein SVM. We have compared the protein encoding/feature extraction from Protein BERT to other sequence-derived feature extraction methods and also compared various ML methods. In addition, genome-wide surveys identified the phyletic distribution of the S9 family members and domain architectures that are prevalent in various phyla. The identification of POP-specific clusters shows that POPs can be sub-classified into 7 different subcategories.

## Results and Discussions

### Identification of POP homologues in Bacterial Lineage

The primary dataset was downloaded from the assembly database of NCBI. (information regarding all the downloaded genomes is available in Additional File 1). The filters used were the availability in latest RefSeq genomes and completeness of assembled genomes. As of Feb 2023, the total number of genomes considered after the above filters is 33018. The distribution of genomes in the dataset shows that ∼45% of the downloaded genomes belong to *Gamma-proteobacteria* followed by *Bacillota*, *Actinomycetes* and *Bacteroidetes* respectively (Additional File 2). Due to these heavy biases in genome distribution, all the calculated properties are normalized by the number of genomes. All the annotated S9 family sequences were then identified. The annotations include S9 family peptidase, Prolyl oligopeptidase family peptidase, DPP IV and Oligopeptidase B. We have observed that certain sequences, annotated as members of the "S9 family," lack the characteristic catalytic domain. These sequences often exhibit similarity to other serine protease families, such as carboxylase, protease II, or the broader superfamily of α/β hydrolases. While they may be homologues to the S9 family, they do not truly belong to the same family. To determine the true members, we carefully analysed the domain architecture of each protein. Only those proteins containing both β-propeller and catalytic domains or at least the catalytic domain are considered as members of the S9 family(Fig 1B). We employ HMMSCAN from HMMER (23) software to identify and filter these domains (24). The results from HMMSCAN were systematically analysed to give each sequence a unique domain architecture (Fig 1B). In total, the initial annotation has 61600 sequences associated with S9 family proteins, involving 32189 genomes. Many of these are 100% identical sequences present in different genomes. They can be referred to as identical homologues or orthologs. The unique number of annotated sequences after data filtering is 31006(Fig 1B). If we include identical homologues(100% sequence identity that are present in multiple species) after considering all filters, the total number of annotated sequences is 52695 (Additional file 3). Table 1 contains the number of annotated S9 family proteins and subfamily classification as per NCBI annotation. As evident from Table 1, a few of the DPP IV (#1294) and Oligopeptidase B(#1217) were classified into subclasses. However, sequences containing the annotation “S9 family peptidase” (#13986) and the annotation “Prolyl Oligopeptidase family peptidase”(#14509) have to be reclassified into a sub-category.

**Table 1.** The number of annotated S9 family proteins and subfamily classification as per NCBI annotation. Table 1 shows the original NCBI annotation on the 31006 unique sequences. Some of the Oligopeptidase B and DPP IV are annotated whereas much of the sequences are annotated as Prolyl oligopeptidase family peptidase or S9 family peptidase which could contain sequences from all subfamilies.

|  |  |
| --- | --- |
| Oligopeptidase B (S9A) | 1217 |
| DPP IV (S9B) | 1294 |
| Prolyl oligopeptidase family peptidase (S9A, S9B, S9C, S9D) | 14509 |
| S9 family peptidase (S9A, S9B, S9C, S9D) | 13986 |
| **Total S9 family proteins** | **31006** |

### Distribution of S9 Family Protein/POP Homologues across different Phyla and Species

The distribution of POP homologues across various phyla is shown in Fig 2A. The number of genomes across phyla in the dataset is highly skewed (Additional file 2). Hence, we have taken the relative frequency as the number of POP homologues found in a phylum normalised for the total number of genomes within the same(Fig. 2A). (Since POP homologues per phyla/species are being considered, 52,695 sequences are considered including the identical homologues). We have analysed the phylum *Proteobacteria* as alpha, beta, gamma and delta owing to a large number of genomes in each category. It is observed that the phylum *Gemmatimonadota* has the highest density of POP homologues. In this phylum although the number of genomes taken for initial analysis is only four, each of the genomes has an average of 19 POP homologues. We also notice that genomes within the phyla *Acidobacteriota*, *Bacteroidetes/Chlorobi*, *Plantomycetota*, *Actinomyocetota*, *Deinococcus-Thermus Alphaproteobacteria*, *Oligoflexia* and *Cyanobacteria* are enriched in POP homologues. The category “others” in Fig 2A refers to all other phyla considered (Additional File 4). In addition, for the phyla *Armatimonadota* and *Calditrichota,* we have only one genome to consider for each but these two genomes have rather high number of POP homologues (12 and 8 respectively, please see Additional file 5).

**Fig 2.**
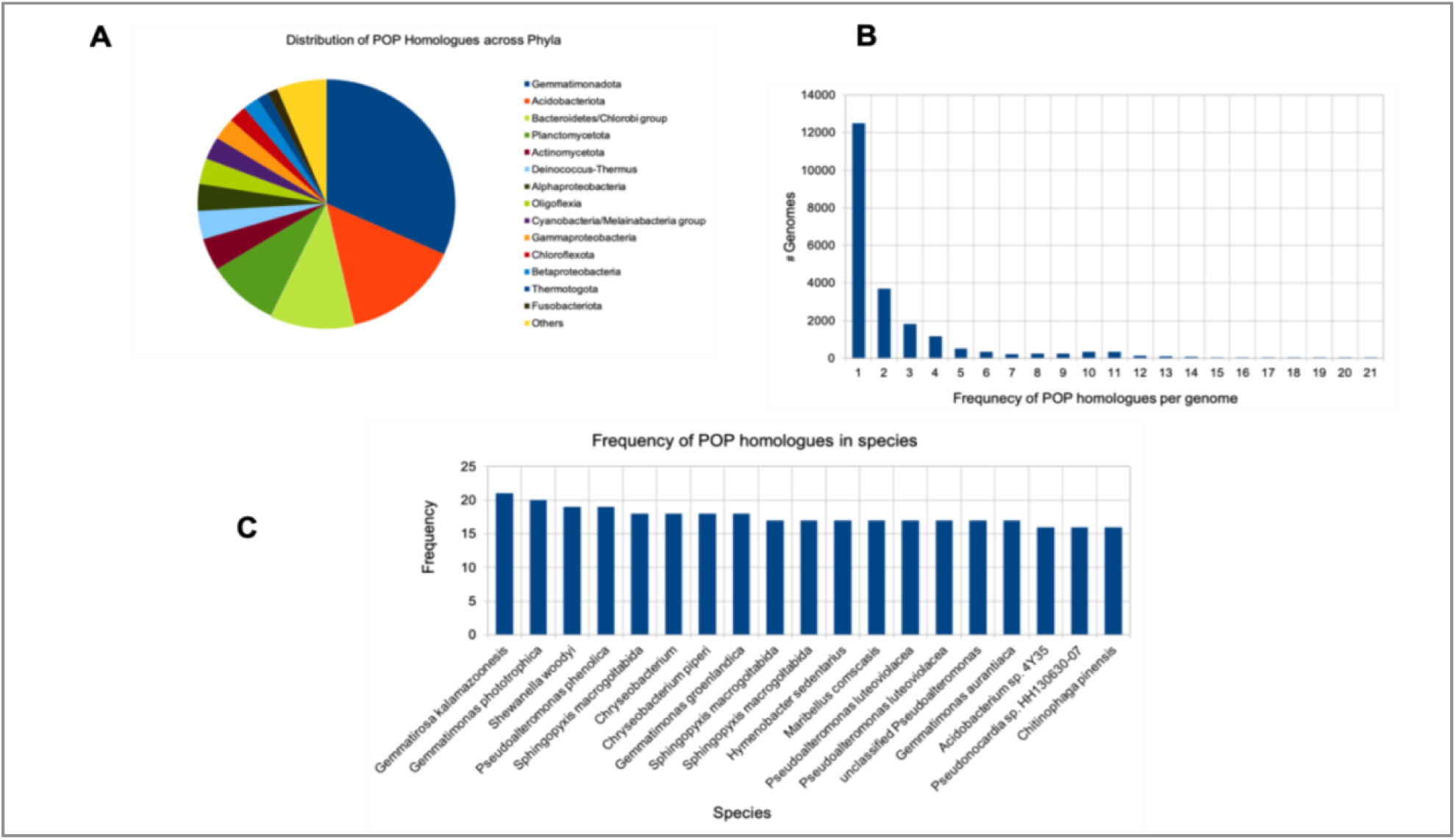
Distribution of POP homologues across phyla and species: The pie chart shows the distribution of POP homologues across phyla. (Fig. 2A). Fig 2B represents the frequency of POP homologues found in individual genomes. Fig 2C shows the species with highest number of POP homologues genes.

Fig 2B shows the histogram of the number of POP homologues across all the species. There is a high variation in the number of annotated POP homologues genes in bacterial genomes, ranging from no POP homologues to multiple copies within a genome. Most of the genomes (#12000) have a single POP family gene. While some of the genomes have POP homologues between 2 and 4, very few species have frequency more than 10. Here the species statistic is calculated by taking individual genome into consideration. *i.e.* considering individual strain (Additional File 6).

Fig 2C shows the distribution of POP homologues in the top 20 highest occurring species/strains. Here, we observe that most of the species with the highest number of POP homologues belong to genus *Gemmatimonas, Shewanella, Sphingopyxis, Chryseobacterium,* and *Pseudomonas*. Most of the genomes have frequency of POP homologues around 18-21, the highest frequency being 21.

### Classifying sub-categories using a Supervised Machine Learning Method

While considering a large number of sequences, we noticed that there is a lot of variation around the active site. Hence, we cannot classify these into subcategories by taking the definition of MEROPS which is based on the fixed motifs around the active site (details in the introduction). No computational method is available that can effectively classify the S9 family proteins in the MEROPs database to the best of our knowledge. Owing to possible cross-talks across subfamilies and a large number of homologues, we have applied a supervised machine learning method for the purpose of classification as labelled examples of subfamily members are available in the MEROPS database.

The training/test set of 960 sequences was derived from the MEROPS examples. Some sequences(particularly for S9B and oligopeptidase B) were also derived from the few annotated sequences of DPP IV and Oligopeptidase B from the analysis. These sequences were randomly selected after a CD-HIT cut-off of 0.8 to not include very similar sequences for training. Additional details including the MEROPS ID of the 960 sequences are added in additional file 7. The catalytic domains of the sequences are used for the training/testing and for predicting the subfamily of new sequences.

Encoding amino acids into numerical values is needed in machine learning (ML) to represent protein sequences as numerical features. The order and composition of amino acids are crucial for understanding the structure and function of the protein of interest. By encoding amino acids into numerical values, ML algorithms can process and analyse protein sequences, enabling various applications in bioinformatics and computational biology (25).

We have used two different methods of encoding: a transformer-based encoding method known as Protein BERT, and sequence-derived features including Amino Acid Composition (AAC), Dipeptide Composition (DPC), and Composition, Transition, and Distribution (CTD).

Protein BERT (26) is a deep language model specifically designed for the analysis of proteins. It is a specialized version of the BERT (Bidirectional Encoder Representations from Transformers) model that has been adapted and fine-tuned for protein sequences. Protein BERT provides an efficient framework for rapidly training protein predictors, even with limited labelled data. To learn the features of proteins, Protein BERT was pre-trained on protein sequences and GO annotations extracted from UniRef90. It also utilizes neighbouring residues context, allowing it to capture local sequence patterns (26). In addition, proteins can vary significantly in length. Transformer-based models, with their self-attention mechanism, can process sequences of variable lengths efficiently (27). The features are also extracted using the AAC, DPC and CTD to compare various feature extraction methods.

The amino acid-encoded feature extraction matrix was derived from both encodings. A split of 70/30 proportion of the dataset was done for training and testing. A parameter grid was set to optimize the hyperparameters along with 5-fold cross-validation for five different ML models namely SVM, Random Forest (RF), Decision tree (DT), K nearest Neighbours (KNN) and Naïve Bayes (NB) (Fig. 3 and additional file 14). This grid encompasses different values for each hyperparameter, allowing for an exhaustive search to identify the optimal combination that maximizes model performance. The details of the hyperparameters optimization, accuracy, balanced accuracy and confusion matrix for each of the models are in the additional file 14. Balanced accuracy was utilized as the metric to evaluate the cross-validation set to determine the most suitable hyperparameters, aiming to address the challenges posed by an imbalanced dataset.

**Fig 3.**
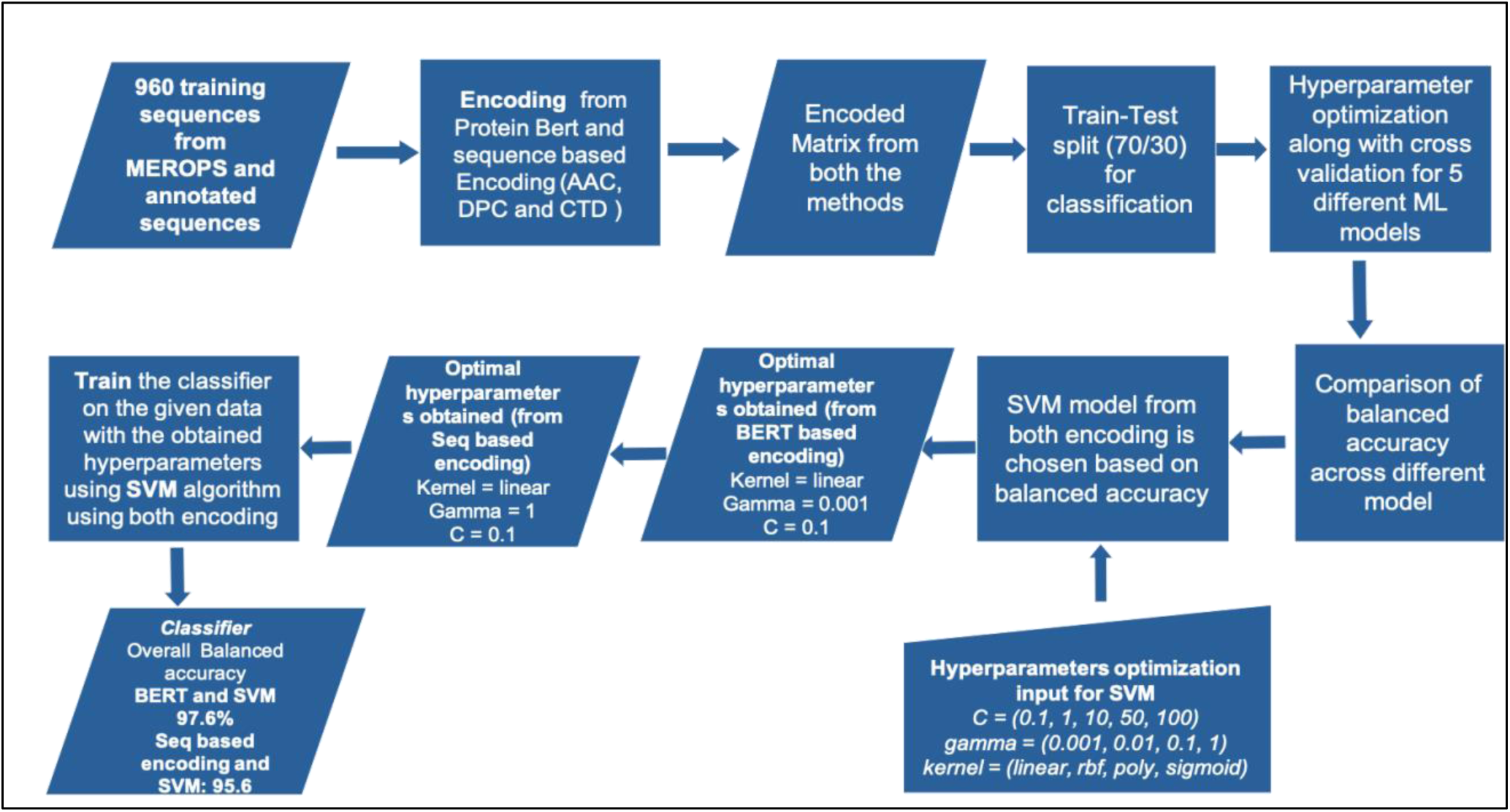
Machine learning (ML) algorithm: Fig. 3 shows the machine learning algorithm used. The five different Machine learning methods used are (SVM, Random forest, Naïve Bayes, Decision trees and KNN). For each model the hyperparameters were optimized using 5-fold cross validation (balanced accuracy is used as the metric for evaluation during cross validation). The balanced accuracy of each of the methods were compared and SVM was found to be better from both the encoding

The SVM model showed the best performance for both encodings, although the performance of Random Forest and KNN was comparable (Fig. 4A). Additionally, for BERT-based encoding, 4 out of 5 ML models achieved better balanced accuracy, though the difference from sequence-based encoding was not significant (Fig 4A).

**Fig 4.**
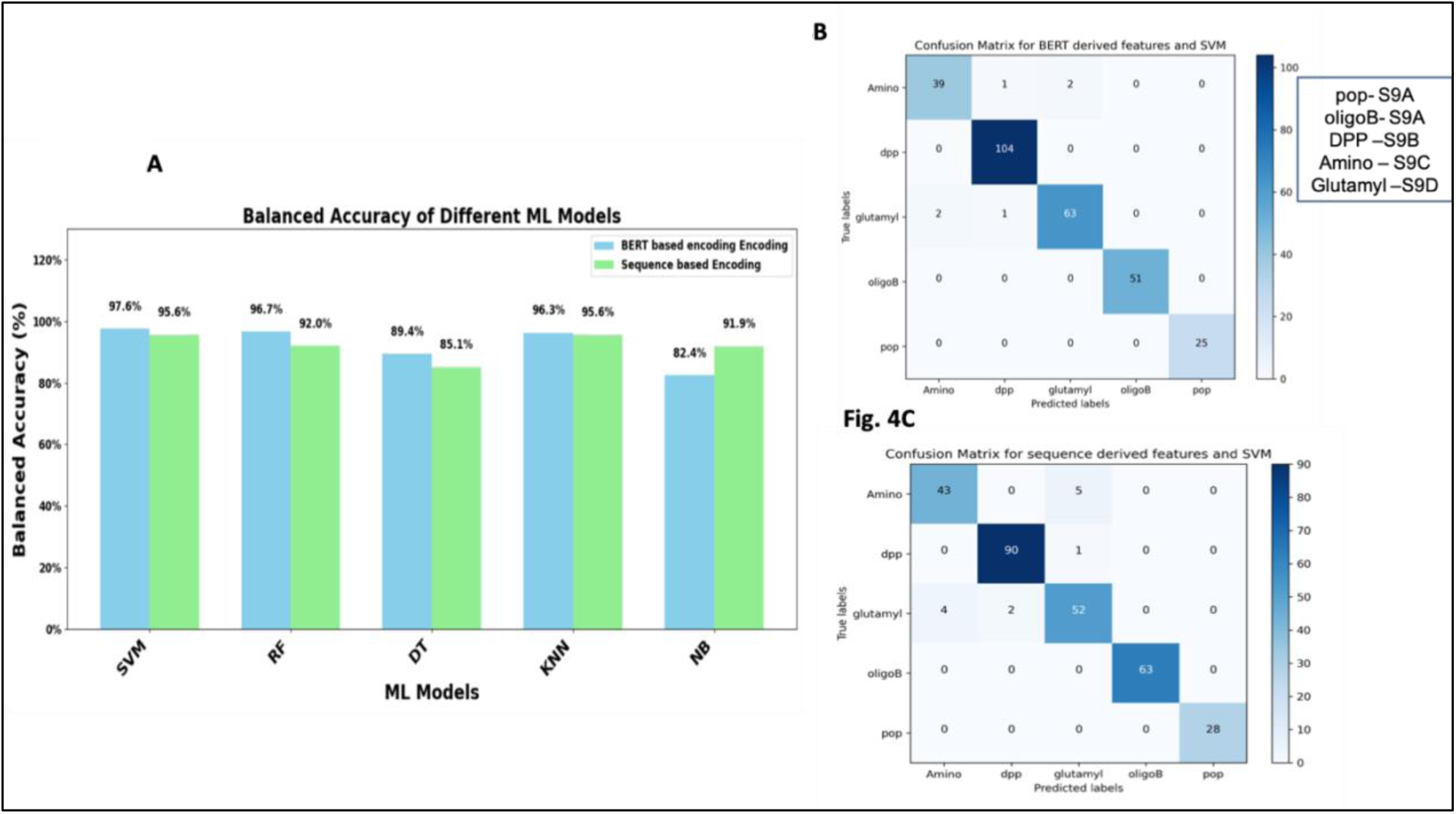
Balanced accuracy of various ML methods and confusion matrix: Fig. 4A shows the balanced accuracy of 5 different machine learning methods for features derived from both BERT and sequence-based methods. The confusion matrix statistics for the SVM classification from both encodings are shown (Fig. 4B and 4C)

The SVM models from both encodings were chosen for the downstream analysis. The optimum hyperparameter obtained for BERT based encoding and SVM was (kernel=’linear’, gamma=0.001, C=0.1) and for Sequence derived encoding and SVM was (kernel=’linear’, gamma=1, C=0.1). SVM is then trained on both the encoded vectors and corresponding labels using the optimized parameters to find the optimal hyperplane that separates different classes.

We have obtained an overall balanced accuracy of 97.6% for BERT based encoding and 95.6 for sequence based encoding. The confusion matrix (obtained from 30% test set after training) (Fig 4B and Fig 4C) shows that all S9A (POP and Oligopeptidase B) are predicted as S9A. While most of the S9B, S9C, and S9D in the test set are predicted accurately, few are predicted in the wrong category. Details of the precision, recall and F1 score for individual categories in tabular format are in Additional file 8.

All other unlabelled S9 peptidases were predicted using these two models for their potential subfamily (results in Additional file 9). Table 2 shows the number of ML method-predicted subfamilies from these two methods and their common predictions. It was noted that more than 90% of the S9A (POPs and Oligopeptidase B) and S9B predictions matched between the two methods, although there were some differences in S9C and S9D. For any downstream analysis, the common sequences from both methods were considered. Overall, approximately 75% of the predictions matched between both methods considering all the category.

**Table 2.** Classification into subfamily of annotated POPs according to the prediction of ML models. Table 2 shows the prediction of subfamilies from both the models. Common sequence from both the models are also shown

| Sub family | Prediction (From BERT and SVM) | Prediction (From seq based encoding and SVM) | Common from Both |
| --- | --- | --- | --- |
| **POP(S9A)** | 5058 | 4767 | 4535 |
| **OPB(S9A)** | 6278 | 6192 | 6057 |
| **DPP IV(S9B)** | 6931 | 5142 | 4497 |
| **Amino acyl peptidase(S9C)** | 5129 | 3846 | 1606 |
| **Glutamyl endopeptidase(S9D)** | 7589 | 11038 | 5836 |

### Additional Validation of ML model

An additional validation was performed to evaluate how the model is performing on unseen data. This is based on how the prediction matches the independent prediction based on domain architecture (not all sequences could be predicted using the domain architecture). Some sequences could be categorized into S9A and S9B from domain architecture if they have the propeller domain. The domain architecture assigns all the catalytic domains from S9A, S9B, S9C, and S9D as S9 catalytic domain. Many sequences have only a catalytic domain or the catalytic domain associated with other domains, they cannot be classified using domain architecture. Also, the domain architecture method cannot classify POP and Oligopeptidase B in the S9A subcategory. It can only classify both of them as S9A. We have independently validated how the prediction method of the ML model agrees with the domain architecture method predictions. Since the S9A and S9B predictions from BERT and sequence based encoding are similar, the predictions from BERT based encoding is considered for this analysis. The number of S9A sequences (POP and Oligopeptidase B) predicted by domain architecture is 10233. The prediction from the ML model as S9A(including predictions for both S9A and S9B) from these 10233 is 10082. The proportion accurately predicted is 98.5%. In addition, the number of S9B(DPP IV) predicted by domain architecture is 3887. The prediction from the ML model as DPP IV from these 3887 is 3857. The proportion accurately predicted is 99.2% (details in additional file 13). Additionally, the ML method is trained only using the catalytic domain. It gives mostly accurate results for proteins that are predicted as S9A and SB based on the domain architecture method, which considers the propeller domain as well. Hence, we believe that it could give reasonably accurate results on categories where the proteins include single catalytic domains, multiple domains with catalytic domains and catalytic domains of S9C and S9D.

### Diverse Domain Architectures (DAs) of Annotated POP homologues

Domain architectures of genes that contain at least one domain in S9 peptidase family were next explored. Frequently occurring domain architectures (with at least 10 representations) are shown in Fig 5A. The most common domain architecture is the single domain one involving only the catalytic domain. This is followed by S9A and S9B family members where the catalytic domain is accompanied by a N-terminal domain. It is important to note that we cannot assign a class based on domain architecture alone as many proteins have only catalytic domains or are associated with other domains like PD40.

**Fig 5.**
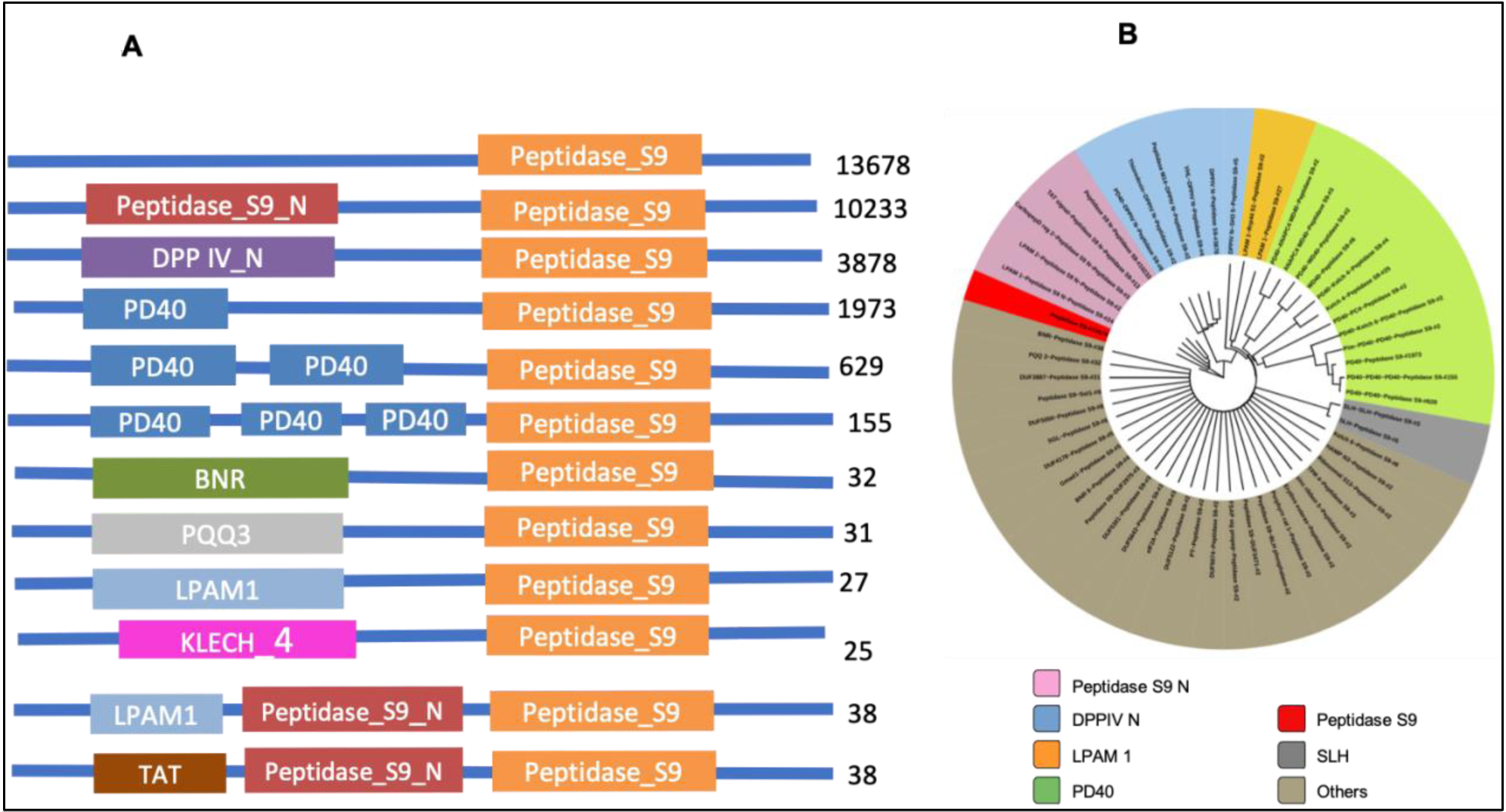
Domain Architecture (DA) representation: Fig 5A represents the domain architecture. The numbers in the right-hand side represents the number pf sequences having that particular DA. ADASS representation of all the domain architecture (Fig 5B). The color coding is done according to the main domain which differentiate the group. The group others represent all single node DAs. They have S9 domains but not associated with combination of other domains

The most common associated (co-existing) domain is PD40 which contains a WD40-like fold. WD40 repeat (WDR) domains are β-propeller-like domains that act as protein interaction scaffolds in multi-protein complexes. Proteins with WD-repeat exhibit high degree of functional diversity and implicated in a variety of functions ranging from signal transduction and transcription regulation to cell cycle control (28). Here we see that β-propeller domain of POP homologues are sometimes replaced with one or more PD40 repeats to accommodate more diverse functions. The other frequently associated domains are PQQ3, LPAM1, Kletch4 and TAT signal. While PQQ, Kletch 4 are also β-propeller like domains, TAT domain is found in translocases which are responsible for the export of folded proteins across the cytoplasmic membrane of bacteria. There are several other domain architectures which are less frequent (found in one or two sequences).

Further, in order to obtain an objective understanding of domain architecture distribution amongst POP homologues, we have applied the ADASS (29) algorithms to show the diversity in domain architectures (Fig 5). Many DAs are represented as a single node as they are not associated with other frequently occurring domains (refer to the colour coding in fig 5B). For example, the DA SGL-PeptidaseS9 is found as a single node, as SGL is not associated with other domains. We also notice that many of the single DA nodes are associated with DUF domains (domains of unknown function). Mostly, the PD40, DPP IV and Peptidase S9_N domains are associated with other domains.

### Distribution of Signal Peptides and Transmembrane Helix across Phyla

Next, we were interested to know the relative abundance of helical regions that can determine the cellular localisation of POP homologues. Most POP homologues from genomes belonging to *Acidobacteriodota*, *Bacteriodetes/Chlorobi* and *Gemmatimonadota* and *Plantomycetota* have signal peptides as well as transmembrane helix(Additional file 16). In addition, putative POP containing genes from the phyla *Armatimonadota* and *Calditrichota* are also enriched in signal peptides and transmembrane helix although they only have single genome representation. Interestingly, although genomes within phyla *Actinomyocetota, Cyanobacteria* and *Bacillota* are enriched in POP homologues, such genes have very few signal peptides and transmembrane helices. We also observe that in the sequences containing S9 domains in bacterial genomes corresponding to the phyla *Nitrospira*, *Oligoflexia,* there are no Transmembrane helix predicted. However, those predicted by us within genomes in the phyla *Spirochaeta, Ternicutes, Fusobacteria* and *Thermotogae* contain neither signal peptides nor transmembrane helix. Detailed information can be found in Additional File 10.

### POP-specific Phylogenetic tree identifies seven distinct clusters/subgroups

Among the members of this S9 family, POPs have garnered extensive research attention. Their capability to cleave a wide range of naturally occurring peptides suggests a regulatory role in processing neuronal peptides and proline-containing hormones (30). Given their substrate specificity and relevance to various human diseases, bacterial pathogenesis, and detoxification processes, these enzymes have become pivotal targets within the pharmaceutical sector. Notably, as illustrated by several studies, both in vitro and in vivo investigations have demonstrated that supplementation with Proline or Glutamine-specific endo-proteases significantly enhances the intrinsic detoxification ability of these enzymes against dietary gluten (31)(32)(33)(34). There could be a potential therapeutic for celiac disease. In addition, they are also implicated in neuronal disorder (15)(35)(36). In light of these significant attributes, our study has undertaken a detailed examination of POPs.

Employing a stringent CD-hit filter (80%) on the common POP predicted data from both the encoding method, we have identified around 1450 representative bacterial POPs for comprehensive analysis, ensuring that the resulting insights accurately reflect the diversity within this enzyme family. A POP-specific phylogenetic tree was constructed and very small clades with less than 30 members are excluded. The POP specific phylogenetic tree represents seven major clusters (Fig 6).

**Fig 6.**
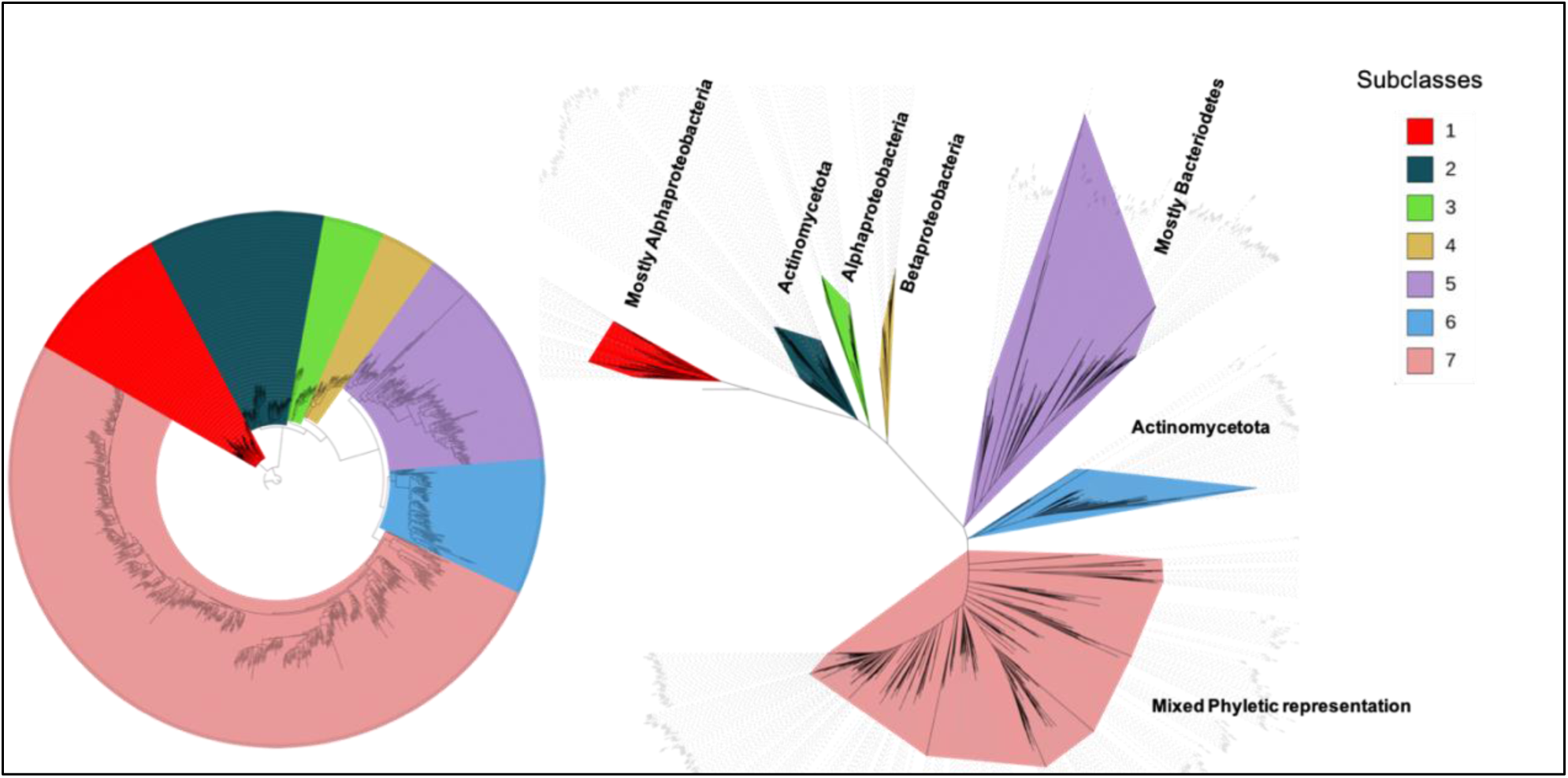
POP specific phylogenetic tree: The POP specific phylogenetic tree (left -circular view, right - unrooted view). Different subclasses with their phyletic representations are shown in different colours

In addition, we also observe that there is co-clustering as well as phyla specific clusters for some phyla across different members of the phyla (Additional file 11 and Fig. 6). Notably the largest cluster (cluster 7) shows co-clustering. This suggests there might be a common function, ancient and conserved across different phyla, much before the phyletic variations emerged.

### Identification of subclass-Specific motifs from the POP-specific phylogenetic tree

We extensively examined these clusters/subclasses to uncover distinct motifs that might underlie their unique characteristics using STREME (37) of MEME (38) tool. Our analysis revealed numerous motifs that are specific to particular classes (Additional File 12). Only those motifs were considered if they were found in more than 95% of the sequences within the cluster. These distinctive motifs play a pivotal role in categorizing the POPs into seven distinct groups. These motifs were found in both catalytic and propeller domains and they are found in both the core of the protein as well as in the surface exposed region (Fig 7). Clusters 2 have the highest number of identified class-specific motifs (31 motifs). Clusters 1 and 6 each have 3 class-specific motifs. Cluster 3 and Cluster 4 each have 10 class-specific motifs. Cluster 5 has 12, and Cluster 7 has 15. Details of motifs for each cluster are in additional File 12 and additional file 15, including the start and end points for each motif in each sequence where it is identified and the background amino acid frequencies.

**Fig 7.**
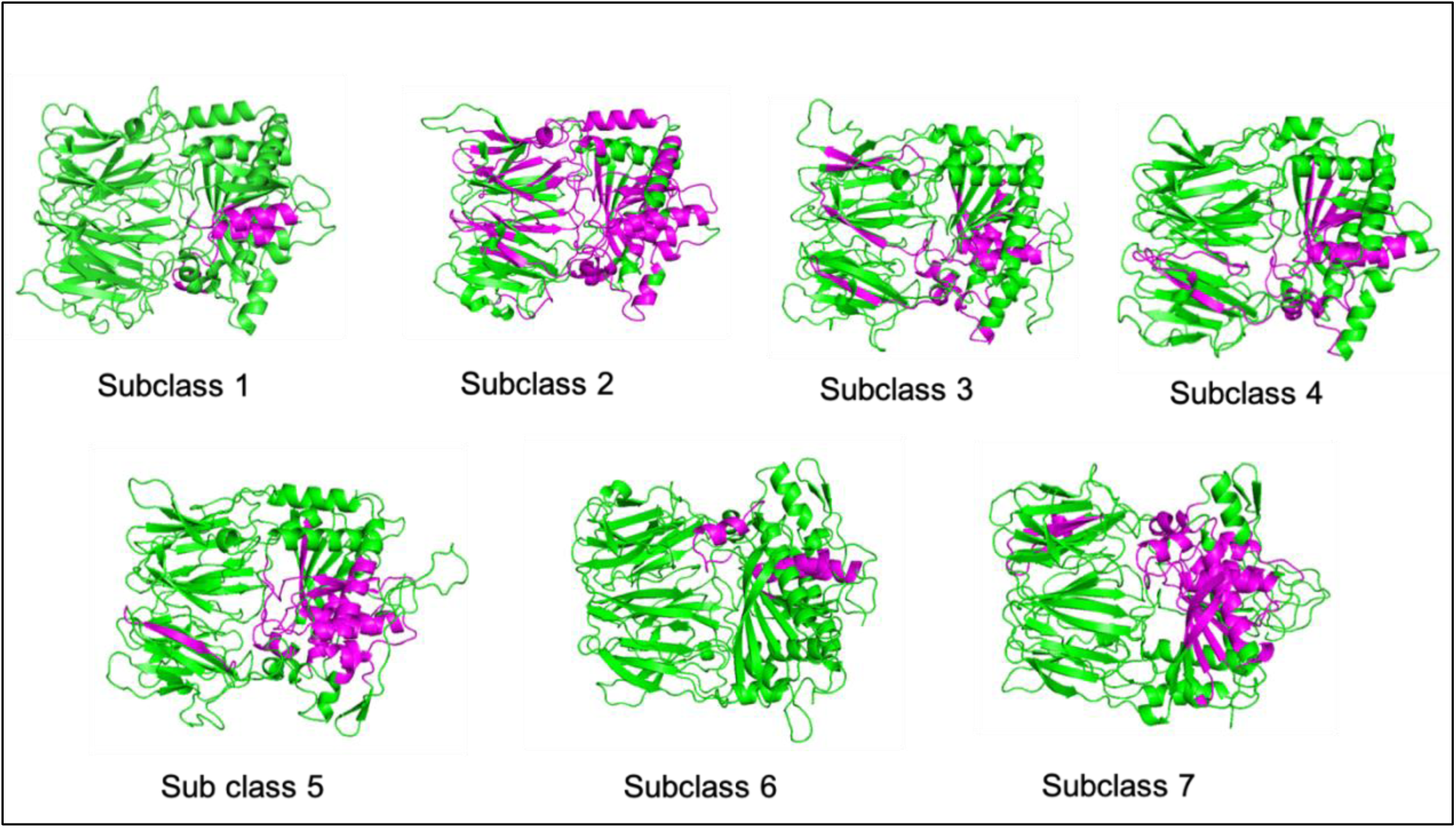
Representation of POP subclass motifs: Fig 7 represents the class-specific motifs positions (in magenta) on representative POP structures from each subclass. Subclass 1 and sub-class 6 have motifs only on the catalytic domain whereas subclasses 2,3,4,5, and 7 have class specific motifs on both the domain

Cluster 7 is highly populated with 570 members, while the other clusters are moderately populated with an average of 100 members (Additional file 11).

In addition, we have noticed that the phyletic distribution is also distinct in different clusters. Cluster 1 and cluster 3 are represented by *alphaproteobacteria* while cluster 2 and cluster 6 are represented by phylum *actinomycetota*. Cluster 5 is represented mostly by the Bacteroidetes/Chlorobi group. Cluster 4 is represented by *betaproteobacteria*. Cluster 7 exhibits mixed phyletic representation which includes POPs from all the included phyla. (details in Additional File 11).

POP sequences within clusters 2, 3, 4, 5 and 7 have class-specific motifs present in both domains (propeller as well as catalytic domains), while members in cluster 1 and cluster 6 have class-specific motifs in only the catalytic domain (Fig 7). Most of the motifs identified are located in the catalytic domain. There were no clusters that have class-specific motifs observed within only the propeller domain suggesting that it is a relatively variable domain.

## Conclusion

In summary, our comprehensive investigation delved into nearly 32,000 bacterial genomes, culminating in the identification of nearly 53,000 S9 family genes within these genomes. We have provided insights into the distribution, classification, and domain architecture of POP homologues across diverse phyla and species through rigorous analysis, We have also classified the POP homologues, using machine various learning methods, as they cannot be classified clearly according to MEROPS definition and according to the difference in their active sites. There is a strong requirement for such automatic tools due to the large numbers handled in this analysis as well as due to the evolutionary divergence, where associations might get increasingly challenging for a few homologues that are in the grey area. The adaptability of machine learning was harnessed to effectively capture the multi-faceted variations within this gene family. The algorithm was able to correctly assign sequences of S9 families into subfamilies with only information on the catalytic domain and no data regarding the entire gene and the coexisting domain during training or testing. Our exploration into domain architecture revealed an array of diversities, and complexity inherent within the structural makeup of these proteins. To address the therapeutic potential of bacterial POPs, we narrowed our focus to the study of POP sequences. Through this dedicated analysis, we identified seven distinct sub-clusters of POPs. Following this, class-specific motifs were identified which suggested that there might be differences in their substrate type and specificity.

The findings of this study contribute significantly to the broader understanding of bacterial S9 family genes. This research lays a foundation for further investigations in the realm of S9 family related therapeutic studies by shedding light on the intricate relationships within this gene family.

## Methods

### Identification of POP homologues

POP homologues were identified directly by searching for annotation in the genomes. Various annotations found in NCBI and used for searching for S9 family proteins are “prolyl oligopeptidase family protein”, “S9 family Peptidase”, “DPP IV “and “Oligopeptidase B. CD-HIT at 100% identity is used to filter make the unique set of sequences. This filter might potentially remove those orthologs (across genomes) which retain 100% sequence identity. However, such a filter is meaningful in the current context and analysis since they are unlikely to yield new information either in phylogeny or in the realisation of class-specific motifs.

### Validation of identified POP homologues

All the annotated POP homologues identified were further validated using HMMSCAN of the HMMER (23) package. HMMSCAN identifies domains in the search sequence against the database of HMM of known domains and we have searched against the PFAM (39) database. Only those sequences that have a catalytic domain (S9_catalytic) were selected for further analysis. We filtered the domains using an independent E-value(iE value) of 0.01 and domain overlaps were resolved based on lower iE value. Additionally, sequences with incomplete domains (model coverage less than 0.7) were systematically removed from our analysis (24). We have seen that many sequences, although annotated as S9 family peptidases, do not have the S9 family domain but domains from similar families like protease II, lipase and carboxypeptidase. Such sequences are also excluded from the analysis.

### Assignment of transmembrane regions, signal peptides and cellular localization

The assignment of the transmembrane helix was done using TMHMM3.0 (40). Analysis of signal peptide is done by SignalP6.0 (41). SignalP software also predicted for Lipo-poly signal peptides but for this analysis, only signal peptides are considered.

### Multiple sequence alignment Phylogenetic tree

Clustal Omega was used for the multiple sequence alignment and trimAl (42) software was used to remove the poorly aligned regions to improve the multiple sequence alignment. A sequence similarity network for further classifying the POP subtypes was done using RaxML (43) online server

### Machine Learning

Tensorflow2.0 (44) was used for protein embedding. In-house Python scripts as well as the Python packages scikit-learn (45) (machine learning library) and pandas (46) were used to deploy the ML model. MatPlotLib (47) Python library was used to calculate subsequent statistics and figures.

### Identification of Sub-clusters and Motifs

POP-specific sub-clusters were identified from the Sequence similarity network. STREME (37) tool within the MEME software suite was used to identify Motifs. Notably, we chose STREME over MEME because, within clusters, the sequence count often surpassed 50. It is worth noting that when sequence numbers exceed this threshold, STREME consistently offers superior outcomes compared to MEME. Clustal Omega (48) was used for the multiple sequence alignment.

### Mapping the motifs into the modelled structure

MODELLER (49) was employed for homology modelling of the POP sequences, where bacterial POP (PDB id: 3ivm) was chosen as a template. The sequences WP_099623704.1, WP_053855468.1, WP_172786573.1, WP_003794441.1, WP_002984378.1, WP_169165980.1, and WP_240542456.1 are used for clusters 1-7 respectively. Clustal Omega was used for multiple sequence alignment. Modelled POP structures were further used for mapping sequence motifs. Figures were generated using Pymol (50).

## List of abbreviations

POP: Prolyl Oligopeptidase
OPB: Oligopeptidase B
DPP IV: Dipeptidyl peptidase-4
HMM: Hidden Markov Model
ML: Machine Learning
iE value: independent E value
DA: Domain Architecture

## Declarations

## Acknowledgements

The authors would like to acknowledge NCBS (TIFR), SERB, IBAB and DBT for infrastructural and other support.

## Funding

This work was supported by PhD fellowship from Department of Biotechnology (DBT). Authors thank NCBS for infrastructural facilities. RS is a J.C. Bose National Fellow (JBR/2021/000006) from the Science and Engineering Research Board, India. RS would also like to thank Bioinformatics Centre Grant funded by the Department of Biotechnology, India (BT/PR40187/BTIS/137/9/2021) and the Institute of Bioinformatics and Applied Biotechnology for the funding through her Mazumdar-Shaw Chair in Computational Biology (IBAB/MSCB/182/2022).

## Author’s contributions

RS designed the experiments and conceived the idea. SN performed the experiments, analysed the data and wrote first draft of the manuscript and RS improved it.

## Availability of data and materials

All the data are available in supplementary material. The data can be browsed and accessed via online web server also(work in progress)

## Competing interests

The authors declare that they have no competing interests.

## Consent for publication

Not applicable

## Ethics approval and consent to participate

Not applicable

## Notes

### Competing Interest Statement

The authors have declared no competing interest.

## References

1. Hedstrom L. Serine Protease Mechanism and Specificity. Chem Rev. 2002 Dec 1;102(12):4501–24.

2. Rawlings ND, Barrett AJ, Thomas PD, Huang X, Bateman A, Finn RD. The MEROPS database of proteolytic enzymes, their substrates and inhibitors in 2017 and a comparison with peptidases in the PANTHER database. Nucleic Acids Res. 2018 Jan 4;46(D1):D624–32.

3. Di Cera E. Serine proteases. IUBMB Life. 2009 May;61(5):510–5.

4. Patel S. A critical review on serine protease: Key immune manipulator and pathology mediator. Allergol Immunopathol (Madr). 2017 Nov;45(6):579–91.

5. Hengge R, Bukau B. Proteolysis in prokaryotes: protein quality control and regulatory principles. Mol Microbiol. 2003 Sep;49(6):1451–62.

6. Tripathi LP, Sowdhamini R. Genome-wide survey of prokaryotic serine proteases: Analysis of distribution and domain architectures of five serine protease families in prokaryotes. BMC Genomics. 2008;9(1):549.

7. Fülöp V, Böcskei Z, Polgár L. Prolyl Oligopeptidase. Cell. 1998 Jul;94(2):161–70.

8. Venäläinen JI, Juvonen RO, Männistö PT. Evolutionary relationships of the prolyl oligopeptidase family enzymes. Eur J Biochem. 2004 Jul;271(13):2705–15.

9. Shan L, Mathews II, Khosla C. Structural and mechanistic analysis of two prolyl endopeptidases: Role of interdomain dynamics in catalysis and specificity. Proc Natl Acad Sci. 2005 Mar 8;102(10):3599–604.

10. Li M, Chen C, Davies DR, Chiu TK. Induced-fit Mechanism for Prolyl Endopeptidase. J Biol Chem. 2010 Jul;285(28):21487–95.

11. MEROPS - the Peptidase Database [Internet]. [cited 2024 Jan 9]. Available from: https://www.ebi.ac.uk/merops/cgi-bin/famsum?family=S9

12. Cunningham DF, O’Connor B. Proline specific peptidases. Biochim Biophys Acta BBA - Protein Struct Mol Enzymol. 1997 Dec;1343(2):160–86.

13. Jones WM, Manning LR, Manning JM. Enzymic cleavage of the blocked amino terminal residues of peptides. Biochem Biophys Res Commun. 1986 Aug;139(1):244–50.

14. Kaukinen K, Lindfors K, Mäki M. Advances in the treatment of coeliac disease: an immunopathogenic perspective. Nat Rev Gastroenterol Hepatol. 2014 Jan;11(1):36–44.

15. Männistö PT, García-Horsman JA. Mechanism of Action of Prolyl Oligopeptidase (PREP) in Degenerative Brain Diseases: Has Peptidase Activity Only a Modulatory Role on the Interactions of PREP with Proteins? Front Aging Neurosci [Internet]. 2017 Feb 14 [cited 2024 Jan 9];9. Available from: http://journal.frontiersin.org/article/10.3389/fnagi.2017.00027/full

16. Motta FN, Azevedo CDS, Neves BP, Araújo CND, Grellier P, Santana JMD, et al. Oligopeptidase B, a missing enzyme in mammals and a potential drug target for trypanosomatid diseases. Biochimie. 2019 Dec;167:207–16.

17. Rea D, Van Elzen R, De Winter H, Van Goethem S, Landuyt B, Luyten W, et al. Crystal structure of Porphyromonas gingivalis dipeptidyl peptidase 4 and structure-activity relationships based on inhibitor profiling. Eur J Med Chem. 2017 Oct;139:482–91.

18. Mentlein R, Gallwitz B, Schmidt WE. Dipeptidyl-peptidase IV hydrolyses gastric inhibitory polypeptide, glucagon-like peptide-1(7–36)amide, peptide histidine methionine and is responsible for their degradation in human serum. Eur J Biochem. 1993 Jun;214(3):829–35.

19. Juillerat-Jeanneret L. Dipeptidyl Peptidase IV and Its Inhibitors: Therapeutics for Type 2 Diabetes and What Else? J Med Chem. 2014 Mar 27;57(6):2197–212.

20. Wang K, Zhang Z, Hang J, Liu J, Guo F, Ding Y, et al. Microbial-host-isozyme analyses reveal microbial DPP4 as a potential antidiabetic target. Science. 2023 Aug 4;381(6657):eadd5787.

21. Brocca S, Ferrari C, Barbiroli A, Pesce A, Lotti M, Nardini M. A bacterial acyl aminoacyl peptidase couples flexibility and stability as a result of cold adaptation. FEBS J. 2016 Dec;283(23):4310–24.

22. Kaushik S, Sowdhamini R. Distribution, classification, domain architectures and evolution of prolyl oligopeptidases in prokaryotic lineages. BMC Genomics. 2014;15(1):985.

23. Eddy SR. Accelerated Profile HMM Searches. Pearson WR, editor. PLoS Comput Biol. 2011 Oct 20;7(10):e1002195.

24. Iyer MS, Joshi AG, Sowdhamini R. Genome-wide survey of remote homologues for protein domain superfamilies of known structure reveals unequal distribution across structural classes. Mol Omics. 2018;14(4):266–80.

25. Yang KK, Wu Z, Bedbrook CN, Arnold FH. Learned protein embeddings for machine learning. Wren J, editor. Bioinformatics. 2018 Aug 1;34(15):2642–8.

26. Brandes N, Ofer D, Peleg Y, Rappoport N, Linial M. ProteinBERT: a universal deep-learning model of protein sequence and function. Martelli PL, editor. Bioinformatics. 2022 Apr 12;38(8):2102–10.

27. Vaswani A, Shazeer N, Parmar N, Uszkoreit J, Jones L, Gomez AN, et al. Attention Is All You Need [Internet]. arXiv; 2023 [cited 2024 Jan 11]. Available from: http://arxiv.org/abs/1706.03762

28. Schapira M, Tyers M, Torrent M, Arrowsmith CH. WD40 repeat domain proteins: a novel target class? Nat Rev Drug Discov. 2017 Nov;16(11):773–86.

29. National Center for Biological Sciences (TIFR), UAS-GKVK Campus, Bellary Road, Bangalore 560 065, India, Syamaladevi DP, Joshi A, Sowdhamini R. An alignment-free domain architecture similarity search (ADASS) algorithm for inferring homology between multi-domain proteins. Bioinformation. 2013 Jun 8;9(10):491–9.

30. García-Horsman JA, Männistö PT, Venäläinen JI. On the role of prolyl oligopeptidase in health and disease. Neuropeptides. 2007 Feb;41(1):1–24.

31. Wei G, Helmerhorst EJ, Darwish G, Blumenkranz G, Schuppan D. Gluten Degrading Enzymes for Treatment of Celiac Disease. Nutrients. 2020 Jul 15;12(7):2095.

32. Osorio CE, Wen N, Mejías JH, Mitchell S, Von Wettstein D, Rustgi S. Directed-Mutagenesis of Flavobacterium meningosepticum Prolyl-Oligopeptidase and a Glutamine-Specific Endopeptidase From Barley. Front Nutr. 2020 Feb 18;7:11.

33. Moreno Amador MDL, Arévalo-Rodríguez M, Durán EM, Martínez Reyes JC, Sousa Martín C. A new microbial gluten-degrading prolyl endopeptidase: Potential application in celiac disease to reduce gluten immunogenic peptides. Sestak K, editor. PLOS ONE. 2019 Jun 27;14(6):e0218346.

34. Kulkarni A, Patel S, Khanna D, Parmar MS. Current pharmacological approaches and potential future therapies for Celiac disease. Eur J Pharmacol. 2021 Oct;909:174434.

35. Eteläinen TS, Silva MC, Uhari-Väänänen JK, De Lorenzo F, Jäntti MH, Cui H, et al. A prolyl oligopeptidase inhibitor reduces tau pathology in cellular models and in mice with tauopathy. Sci Transl Med. 2023 Apr 12;15(691):eabq2915.

36. Svarcbahs R, Julku UH, Norrbacka S, Myöhänen TT. Removal of prolyl oligopeptidase reduces alpha-synuclein toxicity in cells and in vivo. Sci Rep. 2018 Jan 24;8(1):1552.

37. Bailey TL. STREME: accurate and versatile sequence motif discovery. Birol I, editor. Bioinformatics. 2021 Sep 29;37(18):2834–40.

38. Bailey TL, Johnson J, Grant CE, Noble WS. The MEME Suite. Nucleic Acids Res. 2015 Jul 1;43(W1):W39–49.

39. Finn RD, Bateman A, Clements J, Coggill P, Eberhardt RY, Eddy SR, et al. Pfam: the protein families database. Nucleic Acids Res. 2014 Jan;42(D1):D222–30.

40. Krogh A, Larsson B, Von Heijne G, Sonnhammer ELL. Predicting transmembrane protein topology with a hidden markov model: application to complete genomes11Edited by F. Cohen. J Mol Biol. 2001 Jan;305(3):567–80.

41. Teufel F, Almagro Armenteros JJ, Johansen AR, Gíslason MH, Pihl SI, Tsirigos KD, et al. SignalP 6.0 predicts all five types of signal peptides using protein language models. Nat Biotechnol. 2022 Jul;40(7):1023–5.

42. Capella-Gutiérrez S, Silla-Martínez JM, Gabaldón T. trimAl: a tool for automated alignment trimming in large-scale phylogenetic analyses. Bioinformatics. 2009 Aug 1;25(15):1972–3.

43. Stamatakis A. RAxML version 8: a tool for phylogenetic analysis and post-analysis of large phylogenies. Bioinformatics. 2014 May 1;30(9):1312–3.

44. Effective Tensorflow 2 | TensorFlow Core [Internet]. [cited 2024 Jan 12]. Available from: https://www.tensorflow.org/guide/effective_tf2

45. scikit-learn: machine learning in Python — scikit-learn 1.3.2 documentation [Internet]. [cited 2024 Jan 12]. Available from: https://scikit-learn.org/stable/

46. pandas - Python Data Analysis Library [Internet]. [cited 2024 Jan 12]. Available from: https://pandas.pydata.org/

47. Matplotlib — Visualization with Python [Internet]. [cited 2024 Jan 12]. Available from: https://matplotlib.org/

48. Sievers F, Wilm A, Dineen D, Gibson TJ, Karplus K, Li W, et al. Fast, scalable generation of high-quality protein multiple sequence alignments using Clustal Omega. Mol Syst Biol. 2011 Jan;7(1):539.

49. About MODELLER [Internet]. [cited 2024 Jan 12]. Available from: https://salilab.org/modeller/

50. PyMOL | pymol.org [Internet]. [cited 2024 Jan 12]. Available from: https://pymol.org/2/

