## Supplementary material for "Identification and study of Prolyl Oligopeptidases and related sequences in bacterial lineages": Additional_file_8.docx

**Supplementary Table 1: The statistics for the testing set after training for BERT/SVM method.** The statistics of the representative sequences from each class on the test set after the training. The training/testing set was split into a 70/30 pattern for the 960 sequences. 70 per cent of the dataset was used for the training/validation. The table represents the statistics on 30% of the test cases.

| **Sub-family category** | **Precision** | **Recall** | **F1 score** | **Numbers considered in the test set** |
| --- | --- | --- | --- | --- |
| POP(S9A) | 1 | 1 | 1 | 25 |
| Oligopeptidase B(S9A) | 1 | 1 | 1 | 51 |
| DPP IV(S9B) | 0.98 | 1 | 0.99 | 104 |
| Acyl Amioacyl(S9C) | 0.95 | 0.93 | 0.94 | 42 |
| Glutamyl C(S9D) | 0.97 | 0.95 | 0.96 | 66 |

**Overall Test Accuracy: 97.9**

**Overall Balanced Accuracy: 97.66**
