## Supplementary material for "Identification and study of Prolyl Oligopeptidases and related sequences in bacterial lineages": Additional_file_14.docx

**FROM BERT-Based encoding:**

**SVM:**

```
Best Parameters:  {'C': 0.1, 'gamma': 0.001, 'kernel': 'linear'}
Test accuracy: 0.9791666666666666
              precision    recall  f1-score   support

       Amino       0.95      0.93      0.94        42
         dpp       0.98      1.00      0.99       104
    glutamyl       0.97      0.95      0.96        66
      oligoB       1.00      1.00      1.00        51
         pop       1.00      1.00      1.00        25

    accuracy                           0.98       288
   macro avg       0.98      0.98      0.98       288
weighted avg       0.98      0.98      0.98       288

Balanced Accuracy: 0.9766233766233766
Confusion matrix:
 [[ 39   1   2   0   0]
 [  0 104   0   0   0]
 [  2   1  63   0   0]
 [  0   0   0  51   0]
 [  0   0   0   0  25]]
```

**Random Forest**

```
Best Parameters:  {'bootstrap': True, 'max_depth': 10, 'min_samples_leaf': 1, 'min_samples_split': 2, 'n_estimators': 100}
Test accuracy: 0.96875
              precision    recall  f1-score   support

       Amino       0.88      1.00      0.93        42
         dpp       0.97      1.00      0.99       104
    glutamyl       1.00      0.88      0.94        66
      oligoB       1.00      1.00      1.00        51
         pop       1.00      0.96      0.98        25

    accuracy                           0.97       288
   macro avg       0.97      0.97      0.97       288
weighted avg       0.97      0.97      0.97       288

Balanced Accuracy: 0.9677575757575758
Confusion matrix:
 [[ 42   0   0   0   0]
 [  0 104   0   0   0]
 [  5   3  58   0   0]
 [  0   0   0  51   0]
 [  1   0   0   0  24]]
```

**Decision tree**

```
Best Parameters:  {'criterion': 'entropy', 'max_depth': 10, 'min_samples_leaf': 1, 'min_samples_split': 10, 'splitter': 'random'}
Test accuracy: 0.9166666666666666
              precision    recall  f1-score   support

       Amino       0.85      0.83      0.84        42
         dpp       0.93      0.96      0.94       104
    glutamyl       0.89      0.88      0.89        66
      oligoB       0.96      1.00      0.98        51
         pop       0.95      0.80      0.87        25

    accuracy                           0.92       288
   macro avg       0.92      0.89      0.90       288
weighted avg       0.92      0.92      0.92       288

Balanced Accuracy: 0.8947319347319347
Confusion matrix:
 [[ 35   3   4   0   0]
 [  1 100   3   0   0]
 [  4   3  58   0   1]
 [  0   0   0  51   0]
 [  1   2   0   2  20]]
```

**KNN**

```
Best Parameters:  {'metric': 'manhattan', 'n_neighbors': 5, 'weights': 'distance'}
Test accuracy: 0.9756944444444444
              precision    recall  f1-score   support

       Amino       0.95      0.93      0.94        42
         dpp       0.96      1.00      0.98       104
    glutamyl       0.98      0.97      0.98        66
      oligoB       1.00      1.00      1.00        51
         pop       1.00      0.92      0.96        25

    accuracy                           0.98       288
   macro avg       0.98      0.96      0.97       288
weighted avg       0.98      0.98      0.98       288

Balanced Accuracy: 0.9636536796536797
Confusion matrix:
 [[ 39   2   1   0   0]
 [  0 104   0   0   0]
 [  2   0  64   0   0]
 [  0   0   0  51   0]
 [  0   2   0   0  23]]
```

**NB**

```
Test accuracy: 0.8854166666666666
              precision    recall  f1-score   support

       Amino       0.80      0.86      0.83        42
         dpp       0.87      0.96      0.91       104
    glutamyl       0.87      0.92      0.90        66
      oligoB       1.00      0.90      0.95        51
         pop       1.00      0.48      0.65        25

    accuracy                           0.89       288
   macro avg       0.91      0.82      0.85       288
weighted avg       0.89      0.89      0.88       288

Balanced Accuracy: 0.8249769054474936
Confusion matrix:
 [[ 36   2   4   0   0]
 [  2 100   2   0   0]
 [  3   2  61   0   0]
 [  4   0   1  46   0]
 [  0  11   2   0  12]]
```

**From seq-based encoding. With 3 features**

**SVM**

```
Classification Report for Best SVM Model:
               precision    recall  f1-score   support

       Amino       0.91      0.90      0.91        48
         dpp       0.98      0.99      0.98        91
    glutamyl       0.90      0.90      0.90        58
      oligoB       1.00      1.00      1.00        63
         pop       1.00      1.00      1.00        28

    accuracy                           0.96       288
   macro avg       0.96      0.96      0.96       288
weighted avg       0.96      0.96      0.96       288

Fitting 5 folds for each of 1 candidates, totalling 5 fits
Best parameters: {'C': 0.1, 'gamma': 1, 'kernel': 'linear'}

Classification Report for Best SVM Model:
               precision    recall  f1-score   support

       Amino       0.91      0.90      0.91        48
         dpp       0.98      0.99      0.98        91
    glutamyl       0.90      0.90      0.90        58
      oligoB       1.00      1.00      1.00        63
         pop       1.00      1.00      1.00        28

    accuracy                           0.96       288
   macro avg       0.96      0.96      0.96       288
weighted avg       0.96      0.96      0.96       288

Best Parameters: {'C': 0.1, 'gamma': 1, 'kernel': 'linear'}
Test Accuracy: 0.9583333333333334
Balanced Accuracy: 0.9562792092964507
Confusion matrix:
 [[43  0  5  0  0]
 [ 0 90  1  0  0]
 [ 4  2 52  0  0]
 [ 0  0  0 63  0]
 [ 0  0  0  0 28]]
```

**Random forest**

```
Best parameters: {'max_depth': 20, 'max_features': 'sqrt', 'min_samples_leaf': 1, 'min_samples_split': 2, 'n_estimators': 300}
              precision    recall  f1-score   support

       Amino       0.97      0.77      0.86        48
         dpp       0.84      1.00      0.91        91
    glutamyl       0.96      0.86      0.91        58
      oligoB       1.00      1.00      1.00        63
         pop       1.00      0.96      0.98        28

    accuracy                           0.93       288
   macro avg       0.96      0.92      0.93       288
weighted avg       0.94      0.93      0.93       288

Test Accuracy: 0.9305555555555556
Balanced Accuracy: 0.9194376026272579
Confusion matrix:
 [[37  9  2  0  0]
 [ 0 91  0  0  0]
 [ 1  7 50  0  0]
```

**Decision trees**

```
Best parameters: {'criterion': 'gini', 'max_depth': None, 'max_features': None, 'min_samples_leaf': 1, 'min_samples_split': 10}
              precision    recall  f1-score   support

       Amino       0.75      0.81      0.78        48
         dpp       0.89      0.86      0.87        91
    glutamyl       0.68      0.74      0.71        58
      oligoB       1.00      0.95      0.98        63
         pop       1.00      0.89      0.94        28

    accuracy                           0.85       288
   macro avg       0.86      0.85      0.86       288
weighted avg       0.86      0.85      0.85       288

Test Accuracy: 0.8506944444444444
Balanced Accuracy: 0.8512520525451561
Confusion matrix:
 [[39  4  5  0  0]
 [ 1 78 12  0  0]
 [10  5 43  0  0]
 [ 2  1  0 60  0]
 [ 0  0  3  0 25]]
```

**KNN**

```
Best parameters: {'metric': 'manhattan', 'n_neighbors': 3, 'weights': 'distance'}
              precision    recall  f1-score   support

       Amino       0.96      0.90      0.92        48
         dpp       0.96      0.99      0.97        91
    glutamyl       0.93      0.90      0.91        58
      oligoB       0.97      1.00      0.98        63
         pop       1.00      1.00      1.00        28

    accuracy                           0.96       288
   macro avg       0.96      0.96      0.96       288
weighted avg       0.96      0.96      0.96       288

Test Accuracy: 0.9583333333333334
Balanced Accuracy: 0.9562792092964507
Confusion matrix:
 [[43  0  4  1  0]
 [ 0 90  0  1  0]
 [ 2  4 52  0  0]
 [ 0  0  0 63  0]
 [ 0  0  0  0 28]]
```

**Naïve Bayers**

```
Best parameters: {'var_smoothing': 0.43287612810830584}
              precision    recall  f1-score   support

       Amino       0.93      0.88      0.90        48
         dpp       0.85      1.00      0.92        91
    glutamyl       0.93      0.72      0.82        58
      oligoB       1.00      1.00      1.00        63
         pop       1.00      1.00      1.00        28

    accuracy                           0.92       288
   macro avg       0.94      0.92      0.93       288
weighted avg       0.93      0.92      0.92       288

Test Accuracy: 0.9236111111111112
Balanced Accuracy: 0.9198275862068964
Confusion matrix:
 [[42  3  3  0  0]
 [ 0 91  0  0  0]
 [ 3 13 42  0  0]
 [ 0  0  0 63  0]
 [ 0  0  0  0 28]]
```

| Method | BERT BASED | SEQ BASED |
| --- | --- | --- |
| SVM | 97.6 | 95.6 |
| Random forest | 96.7 | 92 |
| Decision tree | 89.4 | 85.1 |
| KNN | 96.3 | 95.6 |
| Naïve Bayes | 82.4 | 91.9 |
