## Supplementary figures and images for "Identification and study of Prolyl Oligopeptidases and related sequences in bacterial lineages"

### Additional file_2.png

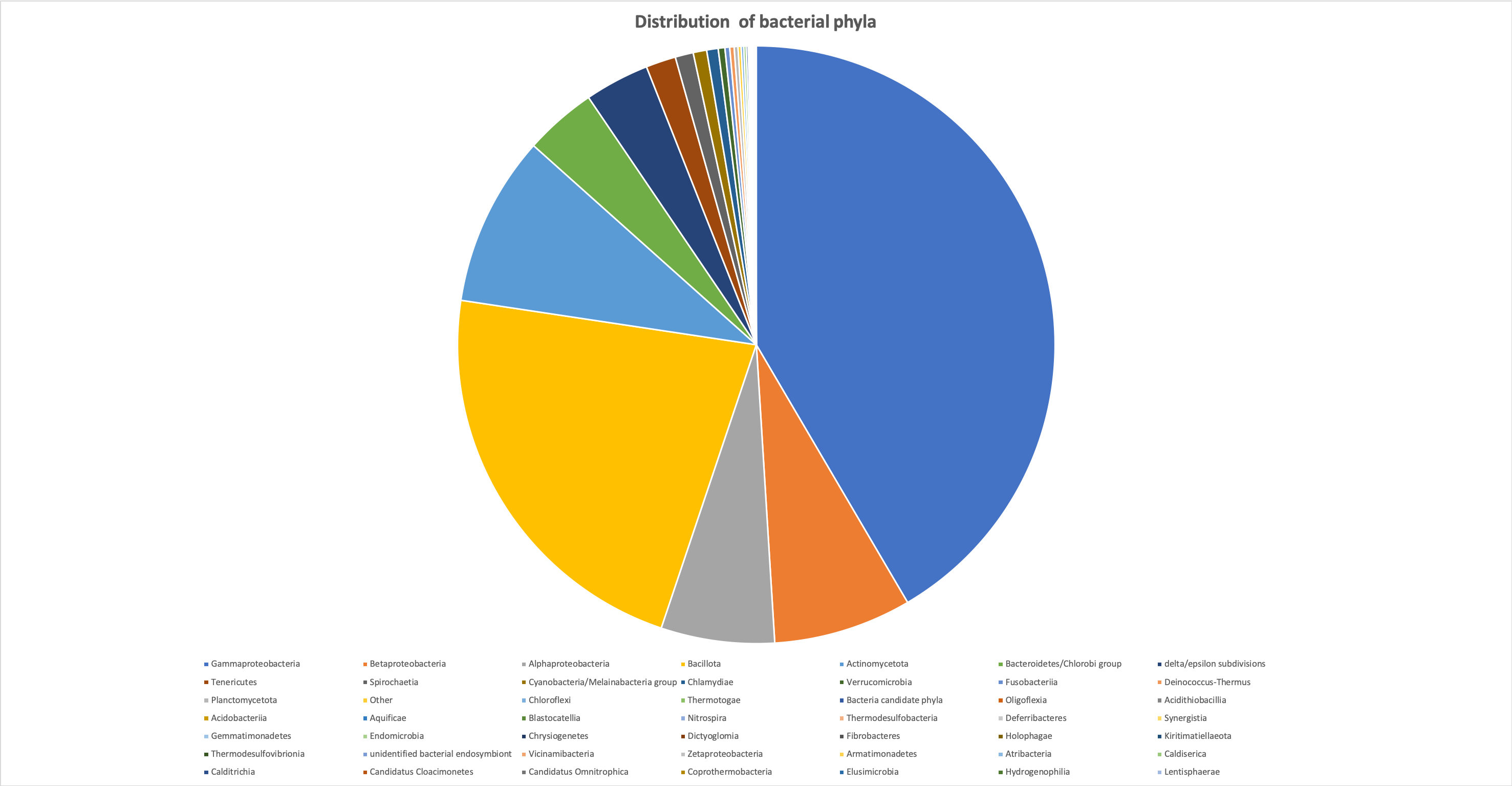

### Additional_file _16.png

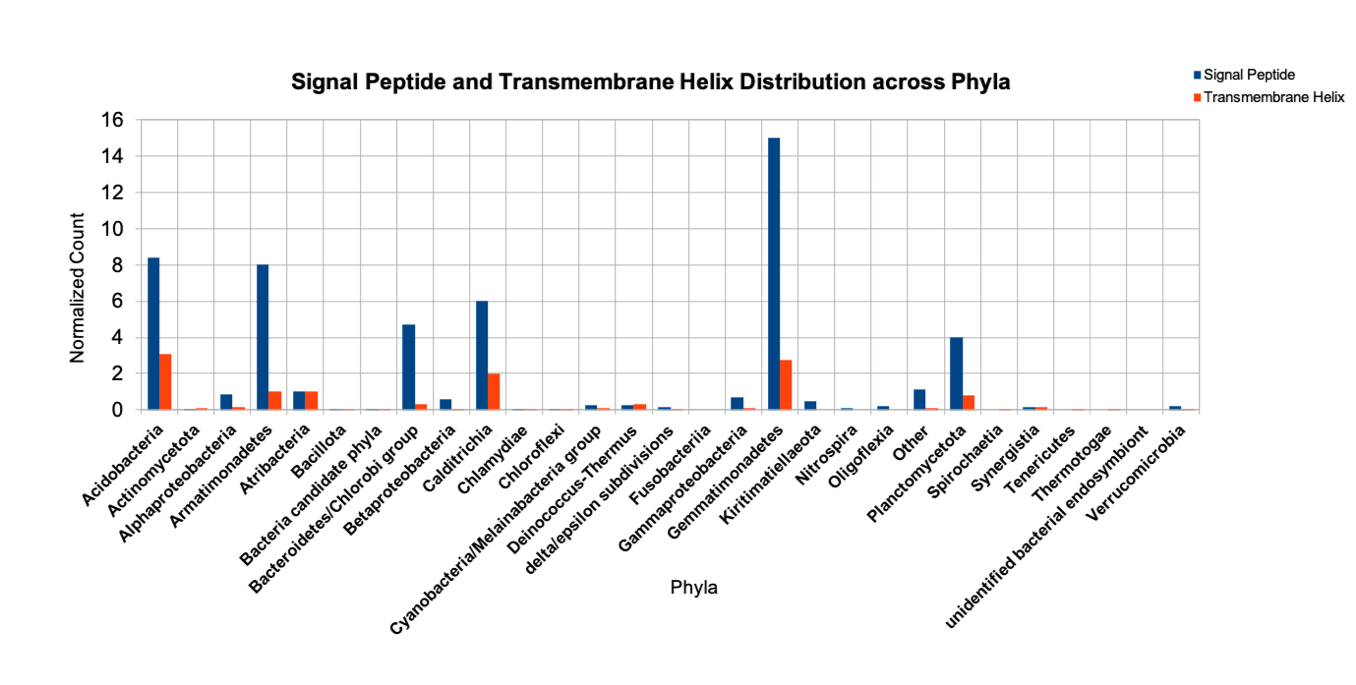
